# IL-22/IL-22RA1 promotes human Tenon’s capsule fibroblasts proliferation and regulates fibrosis through STAT3 signaling pathway

**DOI:** 10.1101/2022.08.09.503275

**Authors:** Yang Zhao, Xiaoyu Zhou, Xinyue Zhang, Dengming Zhou, Baihua Chen, Xuanchu Duan

## Abstract

The study is aimed to investigate that the IL-22/IL-22RA1 signaling pathway regulates scar formation. A total of 31 glaucoma patients who had been previously treated with trabeculectomy surgery and the intraocular pressure was uncontrollable because of scarring and 19 strabismus patients as control patient group. ELISA showed that the IL-22 content of serum from glaucoma patients was 29.80±5.1 ng/μl which is higher than that 5.21±0.9 ng/μl from healthy group significantly. Serum from patients was used to incubate human Tenon’s capsule fibroblasts (HTFs) cells and IL-22 antibody rescued the effect of IL-22 on the biological functions. qPCR and western blot result showed that IL-22 mediates the biological function of HTFs cells via binding IL-22RA1 directly. When transfection of siR-IL-22RA1 or IL-22RA1 gene, the HTFs cells shown significantly anti-fibrosis or pro-fibrosis separately. By using STAT3 inhibitor BAY in IL-22RA1 overexpression group, IL-22-induced proliferation were reduced in HTFS cells. IL-22 promoted fibroblasts cell proliferation and α-SMA via IL-22/IL-22RA1/STAT3 signaling pathway, thereby potentially regulating glaucoma filtration trace fibrosis. This results also show the novel factor in process of postoperative scarring.

**Summary Statement:** The present study suggested that IL-22 expression in glaucoma patient after surgery. IL-22/IL-22RA1 signaling pathway promoted fibroblasts cell proliferation and α-SMA by activating the STAT3 signaling pathway, thereby potentially regulating glaucoma filtration trace fibrosis.

## Introduction

Glaucoma causes blindness accounting for 3.8% of all blind people among age 40 to 80 years in the world(Kang and Tanna, 2021). The number of glaucoma patients over the age 40 has exceeded 9.4 million, of which 5.2 million are monocular blindness and 1.7 million are blindness in both eyes, and the number is expected to reach 111.9 million in 2040(Tham et al., 2014). As the first irreversible blinding eye disease, blindness and disability rates induced by glaucoma have a serious impact on the quality of life of patients. To date, intraocular pressure control remains the mainstay of treatment, and surgery is often considered when medication and laser therapy fail to lower eye drops(Caprioli and Varma, 2011).

The current surgical treatments for glaucoma include: trabeculectomy, aqueous humor drainage implantation and minimally invasive glaucoma surgery (MIGS). In recent years, MIGS surgeries have emerged in an endless stream. However, procedures for internal drainage of aqueous humor to Schlemm’s canal cannot reduce intraocular pressure below the level of episcleral venous pressure(Kasahara and Shoji, 2021); and external drainage procedures require patients to face stent pigment release, pigment obstruction and wound scarring formation(Laroche et al., 2019). From a long-term point of view, trabeculectomy is still a recognized surgical method for lowering intraocular pressure, providing patients with reliable and durable effective intraocular pressure. However, postoperative scarring of the filtering tract, resulting in uncontrolled intraocular pressure and optic nerve damage, is still a vital cause of blindness in glaucoma patients.

Transforming growth factor-β (TGF-β), the main inflammatory factor in the filtration tract, stimulates the proliferation of human Tenon’s capsule fibroblasts (HTFs), and activates cells to be myofibroblasts. Moreover, HTFs cells synthesize and secrete a large amount of extracellular matrix (ECM) to forming wound scar(Zada et al., 2018). In this process, a durable immune response, and the release of inappropriate immune mediators play an important role.

The expression of postoperative immune factors increased, such as (interleukin, IL)-1β, IL-6, IL-8, IL-17, IL-22, etc.(Gajda-Derylo et al., 2019). IL-22 is classified into the IL-10 cytokine family, which is one of the cytokines in immediate response to tissue damage(Wolk and Sabat, 2006). It is mainly secreted by CD4+ T cells, Th cells, and Th22 cells. It is the only known immune factor secreted by immune cells and mainly acts on non-immune cells(Arshad et al., 2020). As a necessary factor for the repair of non-immune cells (epithelial cells, mesenchymal cells, fibroblasts), IL-22 affects the wound healing process, and plays an important biological function after binding to the IL-22 receptor complex, which is a heterodimeric transmembrane receptor composed of interleukin 22 receptor alpha 1 (IL-22RA1) and interleukin 10 receptor 2 (IL-10R2) composition. IL-22 stimulates IL22RA1-expressing cells by increasing the phosphorylation level of Janus protein tyrosine kinase (JAK) /Signal Transducer and Activator of Transcription (STAT) 3, is widely involved in cell proliferation, differentiation, apoptosis, and pro-fibrotic effects(Dong et al., 2021). Previous studies have shown that the expression of IL-22RA1 in the fascia tissue of the filtration tract in patients with glaucoma after surgery is increased(Zhao et al., 2019). However, the specific mechanism of the IL-22 effect on filtration tract scarring is still unclear. Therefore, in order to study filtration tract scarring formation and find new therapeutic targets. It is important to understand the characteristics of IL-22 expression and its binding to receptors and analyze the molecular mechanism of downstream signaling pathways.

## Results

### Expression of IL-22 is elevated in peripheral blood from patients with postsurgical scar

IL-22 content was 29.80±5.1 ng/μl in the serum of glaucoma patients with post-surgical scaring (GP) group was significantly higher than 5.21±0.9 ng/μl in that of the control patients (CP) group (Fig1A) To determine whether IL-22 might play a role in filtration track scar tissue, we analyzed the mRNA and protein expression of IL-22 in that tissue. As shown in Fig 1C, IL-22 protein expression was significantly higher in tissues from patient group than that in the healthy group (P<0.05). And there is similar level in mRNA expression (Fig 1B).

**Figure1.**
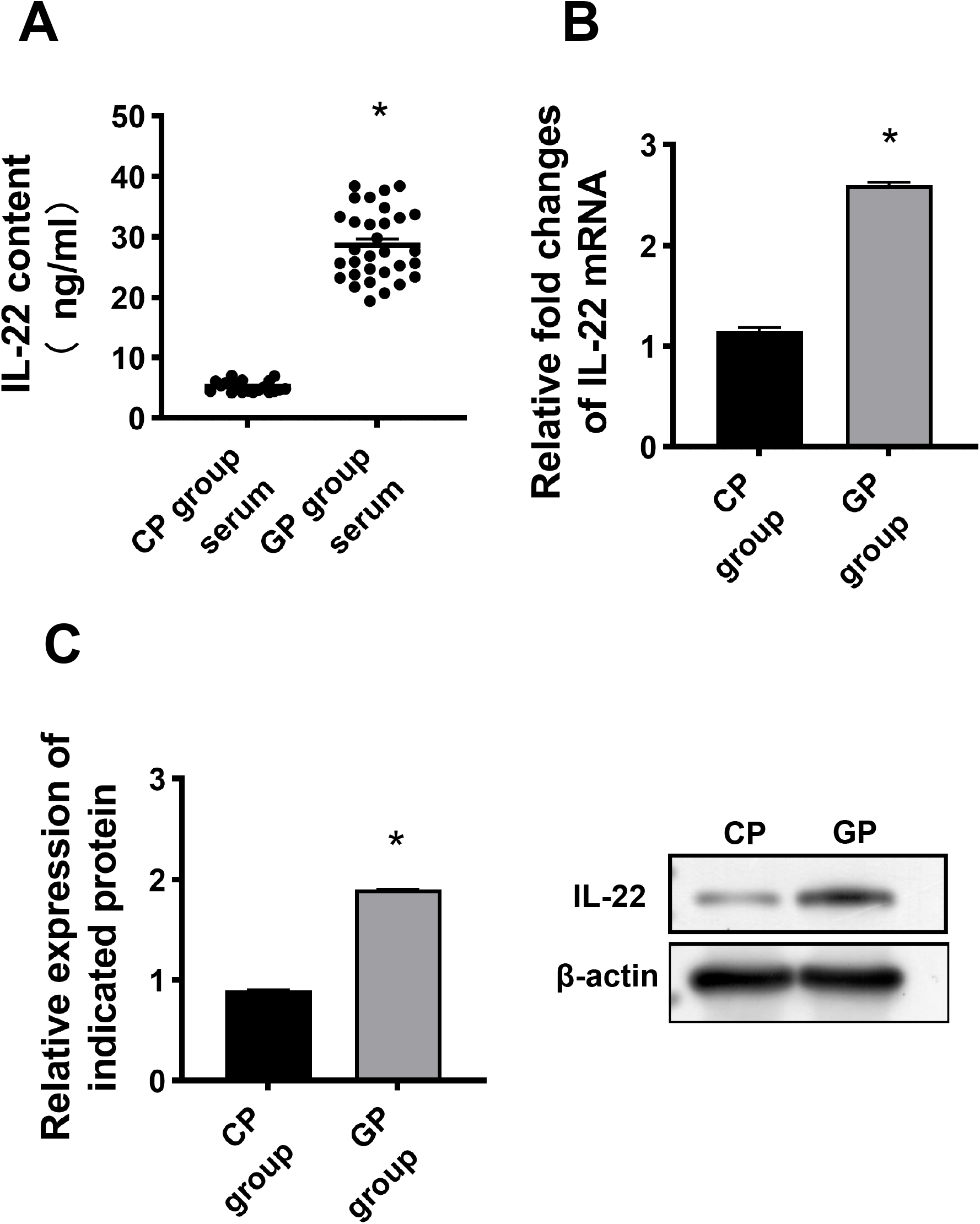
A Content of IL-22 in serum from healthy patients (CP group) and patients with postsurgical scar (GP group), determined by ELISA. *P<0.05, as indicated. B Expression of IL-22 mRNA in tissue in control group and patients with postsurgical scar group. RT-PCR was performed to determine mRNA expression. *P<0.05, as indicated. C Expression of IL-22 protein in tissue in control group and patients with postsurgical scar group. Western blot was performed to determine IL-22 protein expression. *P<0.05, as indicated.

### Expression of IL-22RA1 is elevated in postsurgical scar tissue

The specific and sensitive of IL-22 is depend on IL-22RA1 expression and localization. IL-22RA1 shows a broad tissue distribution, and it was cleared that IL-22RA1 is expressed in the tenon’s capsule fibroblasts in our previous study. To achieve the function, we detected IL-22RA1 expression by PCR and Western blotting, and localized IL-22RA1 protein by immunohistochemical techniques in postsurgical scar tissue. Shown in Fig2A, IL-22RA1 mRNA expression was higher in scar tissue than that in healthy tissue. Western blot also confirmed IL-22RA1 protein expression in scar tissue lysates from anti-glaucoma surgery patients and healthy patients (Fig 2B). Confocal microscopy detected plasma membrane localization of IL-22RA1 in tissue (Fig 2C). α-smooth muscle actin(α-SMA) is a marker of HTFs activation. The result shown in Fig 2 that in postsurgical scar tissue, both mRNA and protein level of α-SMA elevated.

**Figure2.**
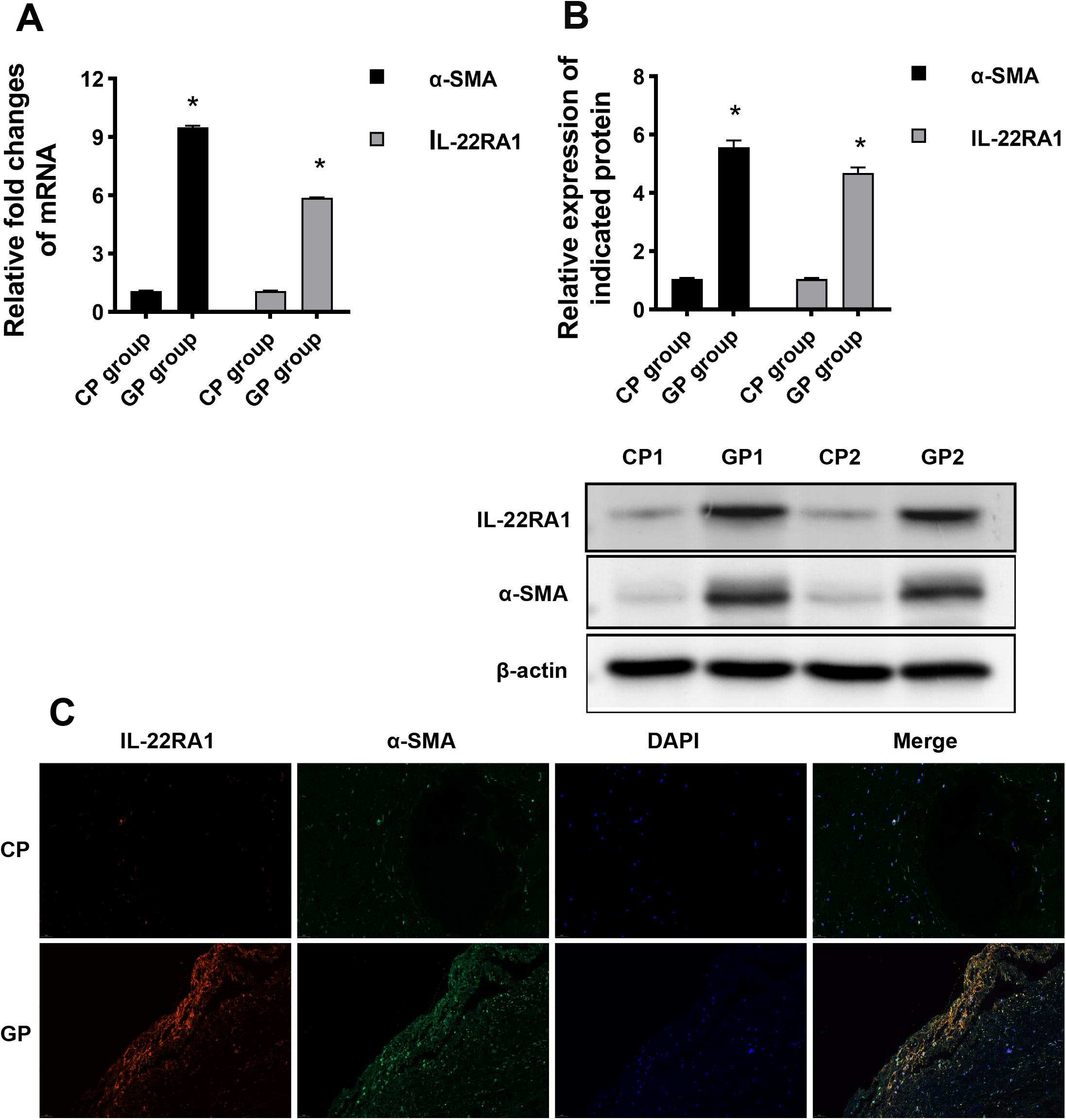
A Expression of IL-22RA1 mRNA in tissue from control group and patients with postsurgical scar. RT-PCR was performed to determine IL-22RA1 and α-SMA mRNA expression. *P<0.05, as indicated. B Expression of IL-22RA1 and α-SMA protein in tissue from control group and patients with postsurgical scar. Western blot was performed to determine protein expression. *P<0.05, as indicated. C IL-22RA1 and α-SMA localization in human tissues. Red: IL-22RA1; Green: α-SMA; Blue: DAPI.

### Glaucoma patient serum stimulates HTFs cell proliferation and activation

After incubation with healthy or glaucoma patient serum for 24 hours, the levels of IL-22RA1 mRNA in HTFs cells were determined by RT-qPCR. The expression of IL-22RA1 mRNA in HTFs cells treated with serum from GP group was significantly higher than that in untreated HTFs cells (CM) (P<0.05), and there was no significant difference between the control group and the healthy serum group (CP group) (Fig 3A). As shown in Fig 3B, the proliferation of HTFs cells at 0, 12, 24, 48 and 72 h after serum treatment was significantly higher than that of untreated (P<0.05), and there was no statistical significance in the control group compared with the healthy serum group (P<0.05). The proportion of G1 phase cells of HTFS cells treated with patient serum was significantly lower than that of the control group (P<0.05; Fig 3C), and there was no significant difference in the proportion of G1 cells between the control group and the healthy serum group. Moreover, the proportion of S-phase cells in HTFS cells treated with serum from glaucoma patients was higher than that in the control group, and there was no significant difference between the control and healthy serum groups.

**Figure3.**
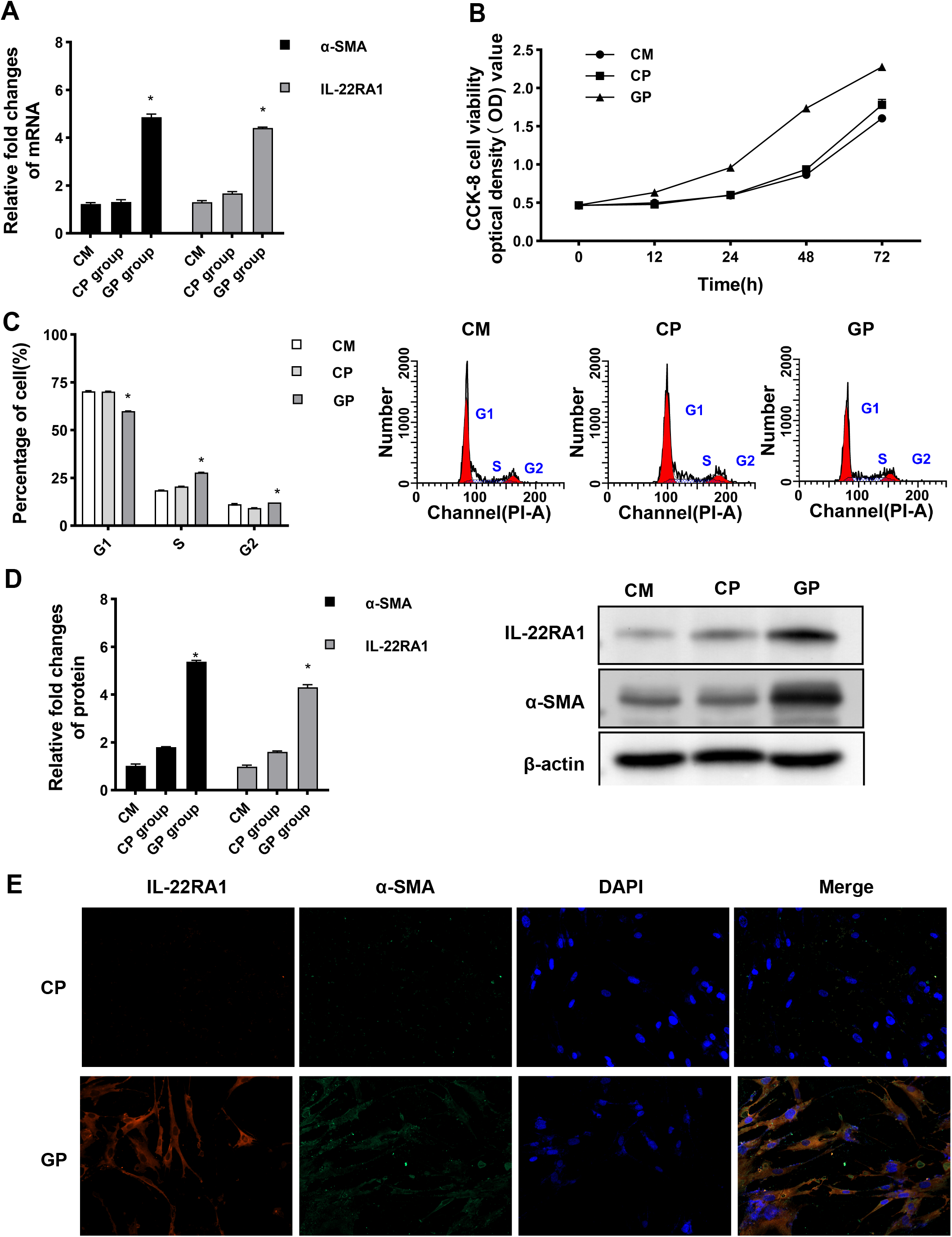
A Expression of IL-22RA1 mRNA in untreated HTFs cells and cells treated with complete medium, healthy or patient serum for 24 h. PCR was performed to determine IL-22RA1 mRNA expression. *P<0.05, as indicated. B Comparison of cell viability was measured by the MTT assay. HTFs cells incubated with complete medium, healthy or patient serum for 12,24,48,72h. All experiments were performed at least three times and results are presented as the means ± SD. *P <0.05, as indicated. C Flow cytometry analysis for cell cycle distribution of HTFs cells incubated with complete medium, healthy or patient serum. The result of one representative assay from three similar independent experiments is shown *P<0.05 vs. Health serum group of the same cell cycle phase. D Expression of IL-22RA1 and α-SMA protein in HTFs cells treated with complete medium, healthy or patient serum. Western blot was performed to determine protein expression. *P<0.05, as indicated. E IL-22RA1 and α-SMA localization in HTF cells. Red: IL-22RA1; Green: α-SMA; Blue: DAPI.

In addition, the patient serum also plays a role in HTFs cell activation. The mRNA and protein expression levels of IL-22RA1, α-SMA were significantly higher than those in the control group (P<0.05; Fig 3A, D), and there was no statistical difference in the protein expression of the control group compared with the healthy serum group.

### IL-22 mediates the biological function of HTFs cells

To further investigate that IL-22 regulates the proliferation and activation of HTF cells, IL-22 antibody were used to conduct rescue experiment. After adding IL-22 antibody, the proliferative capacity of HTF cells was reduced in patients’ serum compared with the health serum group, which was like that in control group (Fig 4A). Furthermore, shown in figure 4B in the IL-22 antibody group cell cycle from the G1 phase to the S phase was inhibited compared to the serum-only group, and reaching the like those in the control group. Moreover, the IL-22RA1 protein expression in the IL-22 antibody group was significantly lower than that in the patients’ serum group (P<0.05; Fig 4C) and like that in control group. In addition, the protein expressions of α-SMA in the IL-22 antibody group were significantly lower than those in the patient’s’ serum treated group (P<0.05; Fig 4C). Taken together, these results suggest that the expression of IL-22RA1 is elevated in HTFs treated with patients’ serum, IL-22 antibody could rescue the IL-22 biology function sot that IL-22 mediated HTFs proliferation and activation directly.

**Figure4.**
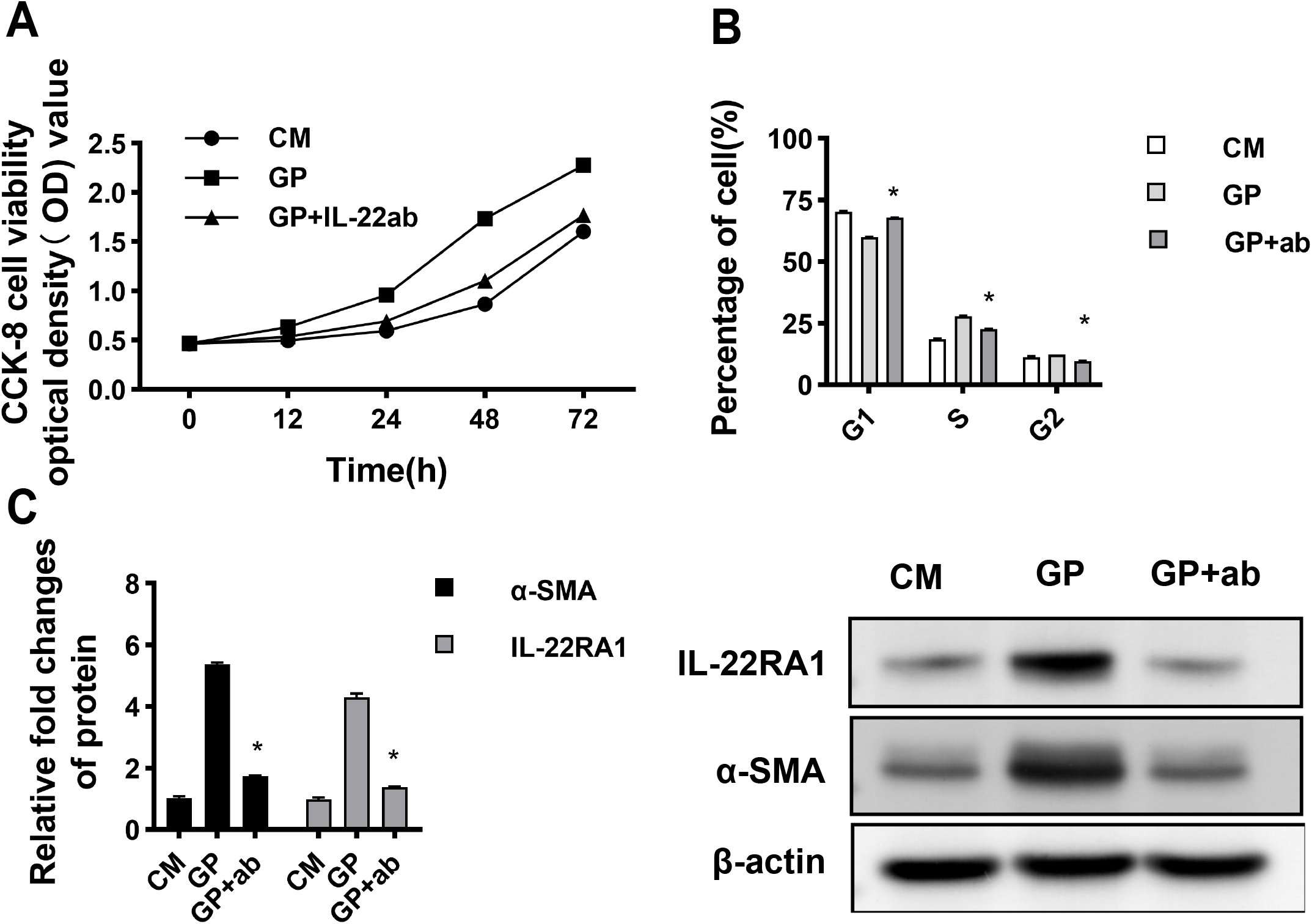
A Comparison of cell viability was measured by the CCK-8 assay. HTFs cells incubated complete medium, in patient serum with/without IL-22 anti-body for 12,24,48,72h. All experiments were performed at least three times and results are presented as the means ± SD. *P <0.05, as indicated. B Flow cytometry analysis for cell cycle distribution of HTFs cells incubated with complete medium, in patient serum with/without IL-22 anti-body. The result of one representative assay from three similar independent experiments is shown *P<0.05 vs. Health serum group of the same cell cycle phase. C Expression of IL-22RA1 and α-SMA protein in HTFs cells treated with complete medium, in patient serum with/without IL-22 anti-body. Western blot was performed to determine protein expression. *P<0.05, as indicated.

### IL-22 exerts its function by binding to IL-22RA1

To test whether IL-22 transmits intracellular signals through IL-22RA1, IL-22RA1 was knocked down or overexpressed in HTFS cells, then subsequently stimulated with IL-22. Fig5A shown the transfection efficiency of siR-IL22RA1 and IL-22RA1. The expressions of IL-22RA1 and α-SMA in the siR-IL-22RA1 group were significantly lower than those in the siR-NC group (P<0.05). In addition, the expressions of IL-22RA1 and α-SMA in the IL-22RA1 overexpression group were significantly higher than those in the NC group (P<0.05; Figure 5B). These results indicated that the transfection experiments were successful. Compared with siR-NC and NC groups, IL-22-induced proliferation was inhibited in siR-IL-22RA1 group, but increased in IL-22RA1 overexpression group. Compared with the siR-NC group and the NC group, the proportion of S-phase cells in the siR-IL-22RA1 group decreased, while the proportion of S-phase cells in the IL-22RA1 overexpression group increased (P<0.05; Figure 5C). The results indicated that IL-22 exerts its biological function through IL-22RA1.

**Figure5.**
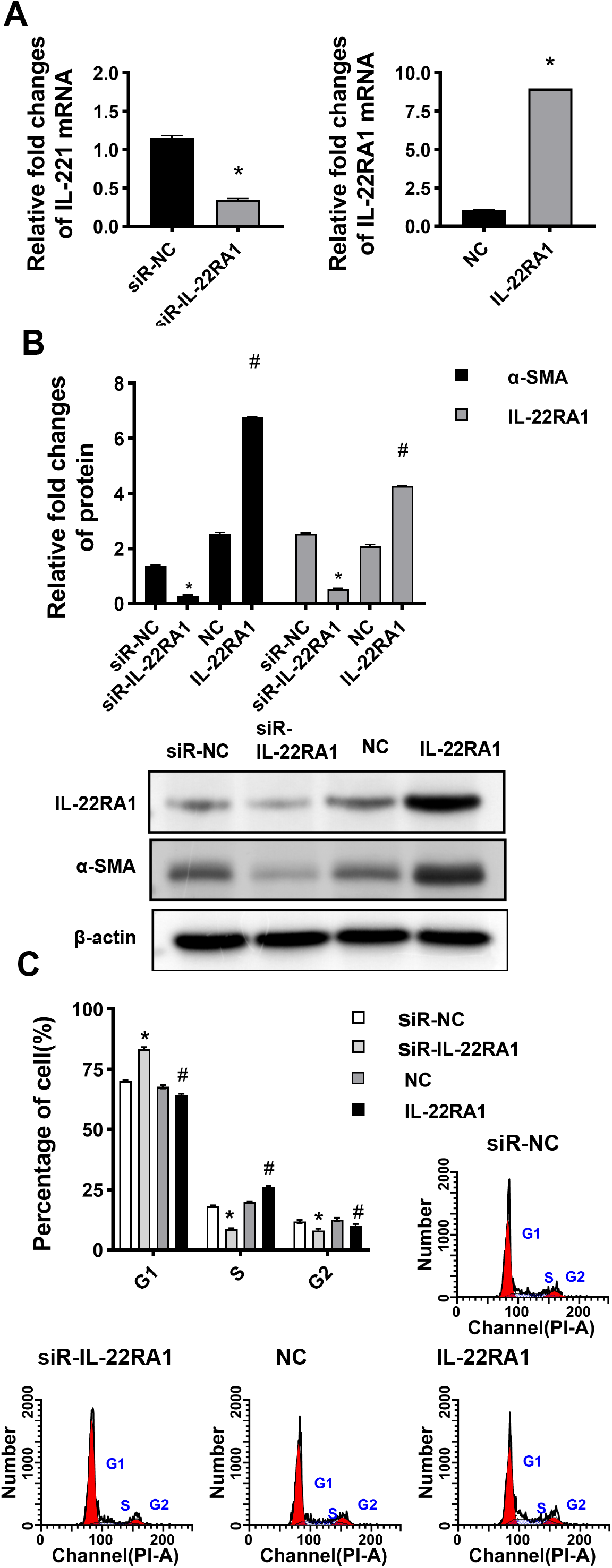
A Transfection efficiency of siR-IL22RA1 and IL-22RA1. *P<0.05 ^*^P<0.05 vs. siR-NC; ^#^P<0.05 vs. NC. B Effect of IL-22RA1 knockdown or overexpression on expression of IL-22R1,α-SMA proteins in HTFs cells transfected with siR-IL-22R1, siR-NC, an IL-22R1 overexpression plasmid or an NC plasmid, incubated with patients’ serum. Western blotting was performed to determine the levels of protein expression. *P<0.05 vs. siR-NC; #P<0.05 vs. NC. C Flow cytometry analysis for cell cycle distribution of HTFs cells transfected with siR-IL-22R1, siR-NC, an IL-22R1 overexpression plasmid or an NC plasmid, incubated with patients’ serum. The result of one representative assay from three similar independent experiments is shown *P<0.05 vs. siR-NC; #P<0.05 vs. NC.

### IL-22/IL-22RA1 regulates the proliferation and activation of HTFs via the STAT3 signaling pathway

To further investigate the mechanism of IL-22RA1 regulates IL-22 functions of fibroblasts, this study investigated the effect of STAT3 signaling pathway inhibitor BAY (2 μM) on HTFS cells. Addition of BAY to the IL-22RA1 overexpression group reduced IL-22-induced proliferation of HTFS cells (P<0.05; Figure 6A). Moreover, the addition of BAY limited the effect of IL-22 on α-SMA expression (P<0.05; Figure 6B). From flow cytometry analysis, the results showed that cells treated with BAY reduced the effect of IL-22 on the proportion of cells transitioning from G1 to S phase (P<0.05; Figure 6C). Overall, these results suggest that IL-22 may regulate the proliferation and expression of α-SMA proteins in HTFS cells via the STAT3 signaling pathway.

**Figure6.**
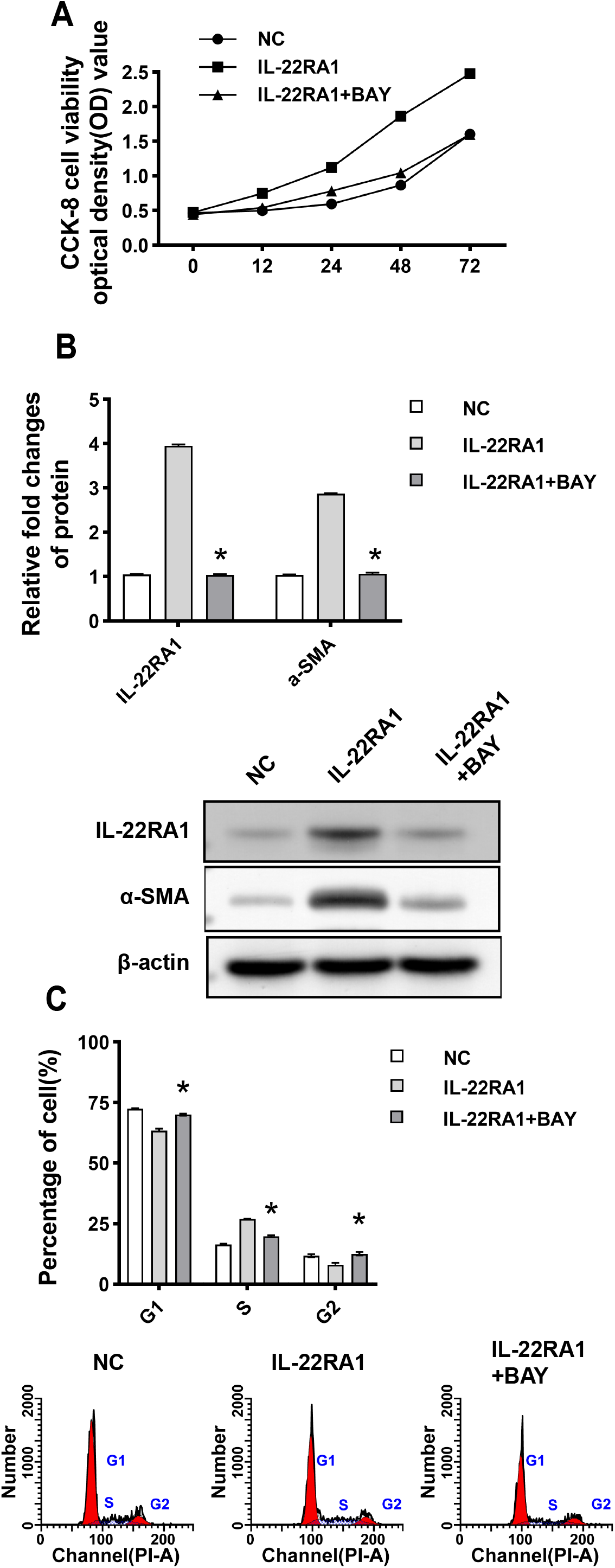
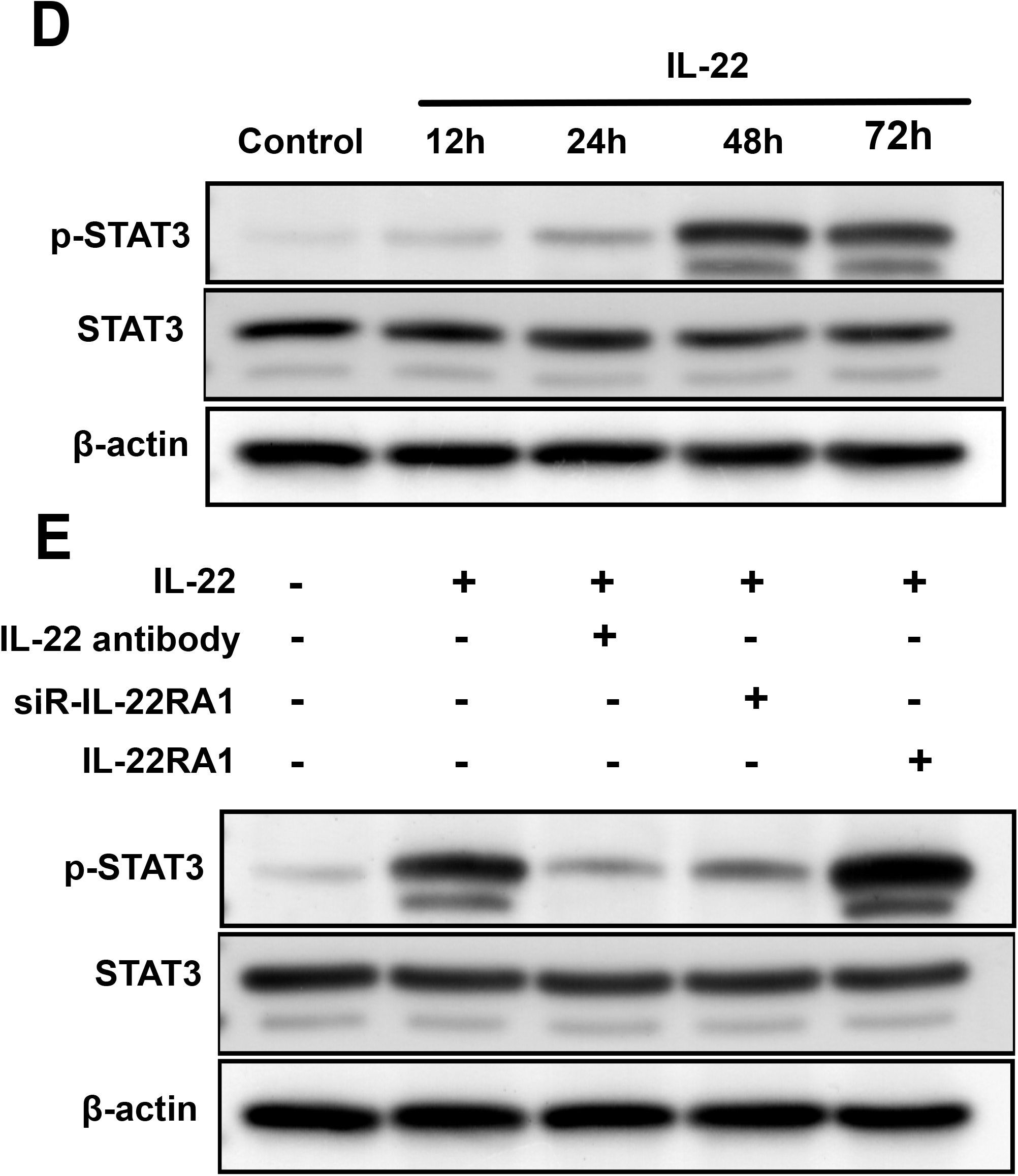
A Comparison of cell viability was measured by the CCK-8 assay. HTFs cells were transfected with an IL-22R1 overexpression plasmid with/without BAY, an NC plasmid, incubated with patients’ serum for 12,24,48,72h. All experiments were performed at least three times and results are presented as the means ± SD. B Expression of IL-22RA1 and α-SMA protein in HTFs cells transfected with an IL-22R1 overexpression plasmid with/without BAY, an NC plasmid, incubated with patients’ serum. Western blot was performed to determine protein expression. *P<0.05, as indicated. C Flow cytometry analysis for cell cycle distribution in HTFs cells transfected with an IL-22R1 overexpression plasmid with/without BAY, an NC plasmid, The result of one representative assay from three similar independent experiments is shown *P<0.05 vs. siR-NC; #P<0.05 vs. NC. D STAT3 phosphorylation induced by IL-22 in HTFs. Cells were cultured with IL-22 for 12, 24, 48, and 72 h. P-STAT3 and STAT3 protein levels were assessed by Western blot analysis. E HTFs cells were stimulated or not for 30 min with IL-22HTFs. Then, cells were incubated with either anti-IL-22RA1 antibody, or transfected with siR-IL-22RA1, IL-22RA1. P-STAT3 and STAT3 protein levels were assessed by Western blot analysis.

Furthermore, IL-22 induced STAT3 phosphorylation in HTFs cells. A time course study showed that IL-22-induced STAT3 phosphorylation increased over time, but from the 72h time point, STAT3 phosphorylation was seen to decrease (Fig. 6D). To further show the role of IL-22RA1 in IL-22 signaling, we investigated the effect of IL-22-induced STAT3 phosphorylation using anti-IL-22 antibody, siR-IL-22RA1 and IL-22RA1. As shown in Figure 6E, HTFs were inhibited IL-22-induced STAT3 phosphorylation by 56% and 83% under anti-IL-22 antibody and siR-IL-22RA1 treatments, respectively. Phosphorylation of STAT3 was increased under IL-22RA1 transfection.

## Discussion

Several factors play a role in filter tract scarring, including the proliferation of fibroblasts, the imbalance of ECM synthesis and degradability, as well as cytokine production(Gajda-Derylo et al., 2019). In various tissues, IL-22 stimulates inflammatory responses, wound healing, and tissue regeneration(Saxton et al., 2021). Previous studies suggest that acinar cells of the lacrimal gland in dry eye mice secrete IL-22, which inhibits IL-17-mediated inflammation of the ocular surface(Ji et al., 2017). Th22 cells, αβTh, and γΔT cells are widely present in the conjunctiva, and IL-22 production benefits the healing of ocular surface wounds. By applying an anti-IL-22 antibody in a mouse corneal epithelial injury model, corneal cell divisions can be significantly reduced by 52%(Yoon et al., 2018). In addition, the expression of IL-22 is upregulated in the inflammatory phase of wound healing in IL-22 knockout mice whose wounds were undergoing abnormal granulation tissue formation, extracellular matrix protein production, and delayed skin wound healing(McGee et al., 2013). There is evidence that IL-22 plays a role in wound healing and tissue regeneration.

Glaucoma patients’ HTFs cells express high levels of IL-22RA1 and this expression increases with scarring. In fibroblasts, IL-22 binds to IL-22RA1, activates downstream STAT3 phosphorylation, and triggers the expression of fibronectin and collagen. In psoriatic dermal fibroblasts, IL-22 binds to IL-22RA1 and stimulates the activation of the JAK/STAT pathway, resulting in increased production of extracellular matrix, fibronectin, and collagen(Zheng et al., 2007). Both IL-20 and IL-22 are members of the IL-10 family. In trabecular cells of glaucoma patients, IL-20 binds to the receptor IL-20 receptor A and participates in extracellular matrix remodeling through the JAK/STAT signaling pathway, leading to increased intraocular pressure(Keller et al., 2014).

In this study, STAT3 inhibitors blocked the IL-22/IL-22RA1 signaling pathway’s effect on HTFs cell proliferation and a-SMA production. HTFs cells induced STAT3 phosphorylation in response to IL-22, thereby directly linking the function of IL-22RA1. Furthermore, both overexpression and blockade experiments demonstrated the requirement for IL-22RA1 for STAT3 phosphorylation. These results suggest that IL-22/IL-22RA1 may promote fibroblast proliferation and activation through the STAT3 signaling pathway in the process of post-surgery filtration tract tissue fibrosis. The accurate mechanism is not completely understood and requires further investigation.

Due to high postoperative fibrosis and failure rates of glaucoma filtration surgery, new drugs and treatments continue to be of interest to improve glaucoma surgical patient outcomes. Rare information reported about IL-22 in glaucoma, so that this study protrudes novel direction anti-fibrosis mechanism after surgery. Due to secretion of IL-22 detectable in serum, in the further IL-22 could be considered as a marker to protect and instruct postoperative treatment.

## Materials and methods

### Patient and tissue collection

Serum and fascia tissue were collected from 31 glaucoma patients (GP group) and 19 strabismus patients as control patient group (CP group) from November 2021 to March 2022 in Changsha Aier Eye Hospital. The criteria for selecting patients were: GP group included glaucoma patients who treated with trabeculectomy surgery already and due to scar formation, the intraocular pressure was uncontrolled; CP group included strabismus patients without other ocular diseases; all patients with no immune diseases, for example ulcerative colitis, atopic dermatitis and rheumatoid arthritis, no recent history of acute infection in the past month. Peripheral blood (5 ml) of all the patients was collected from median cubital veins. By centrifuging at 600 x g and 4°C for 5 min, serum was separated for ELISA test and incubating cells. The fascia tissue samples were snap-frozen in liquid nitrogen and then stored at −80 °C for further use. All procedures conducted in this study were performed in compliance with rules described in the Declaration of Helsinki. All patients provided their written were informed consent prior to surgery and tissue collection. The study protocol was approved by the Human Ethics Committee of Changsha Aier Eye Hospital. (Changsha, China)

### Measurement of serum cytokine level

The level of IL-22 in the patients’ serum were measured by ELISA kits (Quantikine ELISA kits, R&D Systems; Minneapolis, MN, USA). The absorbance of each well was measured with a microplate reader (Multiskan FC, Thermo Fisher Scientific, Waltham, MA, USA) set to 450 nm, with the wavelength correction set to 540 nm.

### Cells culture

HTFs were isolated from individuals undergoing strabismus surgery who had no history of conjunctival disease or use of topical ocular medication as previously described(Zhao et al., 2019). The HTFs were cultured in Dulbecco’s Modified Eagle Medium supplemented with 10% (v/v) fetal bovine serum (Hyclone Laboratories, Logan, UT, USA) and 1% streptomycin-penicillin (Gibco, Thermo Fisher Scientific, Waltham, MA, USA) in a humidified atmosphere at 37 °C with 5% CO_2_. HTFs cultured in above medium says as complete medium group (CM group). Passages 4–6 were used for further experiments. Prior to each experiment, the cells were allowed to reach a sub-confluent status (∼80% confluence); after which, they were cultured in serum-free medium for 24 h.

To investigate the effect of patient serum on HTFs, serum from GP group or CP group was mixed with medium at a ratio of 1:10 to treat HTFs cells. For IL-22 experiments, cells were exposed to different dose-course or time-course of human IL-22 (Peprotech, Rocky Hill, NJ, USA). In order to study the rescue effecting, cells were treated with human IL-22 antibody (1:800, ab133545, Abcam, Cambridge, United Kingdom).

### Cell Transfection

The nucleotide sequences that encoded for IL22RA1 were amplified via PCR performed with the following primers: F, 5′-GGGGTACCATGAGGACGCTGCTGACCATCTTGA-3′; R, 5′-CCGCTCGAGTCAGGACTCCCACTGCACAGTCAGG-3′. Those sequences were then subcloned into the KpnI and XhoI sites of a pcDNA 3.0 vector. An empty pcDNA 3.0 vector was used as a control. siRNA-NC, siR-IL-22RA1 were obtained from GenePharma (Suzhou, China). HTFs were seeded into the wells of six-well plates and cultured to ∼80% confluence. Transfections were performed using Lipofectamine 3000 (Thermo Fisher Scientific, Waltham, MA, USA) as described in the manufacturer’s instructions.

### Flow cytometry

According to the manufacturer’s manual, flow cytometry performed cell cycle analysis. First, preparation cell with DNA Reagent kit (BD Biosciences, San Jose, CA, USA). In briefly, the harvested cells were incubated in sequence with solutions. Then, the cells were analyzed using a flow cytometer and ModFit software (v3.2; Verity Software House, Inc.). Each experiment was repeated three times.

### Cell viability assay

Cell viability was measured by a Cell Counting Kit-8 (CCK8) (Lianke Bio, Hangzhou, Zhejiang, China) according to manufacturer’s instructions. In brief, cells were seeded at a density of 1×10^4^ cells/well in a 96-well plate triplicated. After the cells with specified treatment, a CCK8 working reagent was gently added to each well, and then placed in a 37 °C incubator for 2 h. Finally, the absorbance of each well at 450 nm was measured with a microplate reader (Multiskan FC, Thermo Fisher Scientific, Waltham, MA, USA).

### RNA extraction and PCR analysis

RNA was extracted from HTFs cells using Trizol reagent (Invitrogen, Thermo Fisher Scientific, Waltham, MA, USA). PowerUP SYBR Green Master Mix kit (Thermo Fisher Scientific, Waltham, MA, USA) was used to detect mRNA expression of IL-22 (F 5′-CAACAGGCTAAGCACATGTCATATT-3′; R 5′-TCTCTCCACTCTCTCCAAGCTTT-3′), IL-22RA1 (F 5′-CCTGATGTGACCTGTATCTCCAA-3′, R 5′-GGTCAGGcCGAAGAACTCATATT-3′), α-SMA (F 5′-GAGACCACCTACAACAGCATCAT-3′, R 5′-GCCGATCCACACCGAGTATTT-3′) and GAPDH (F 5′- GGAGTCCACTGGCGTCTTCA-3′, R 5′-GTCATGAGTCCTTCCACGATACC -3′) according to the manufacturer’s protocol. Resulting data were analyzed using the comparative cycle threshold (Ct) method. The target gene cycle thresholds were adjusted relative to a calibrator (normalized Ct value obtained from control groups) and expressed as 2−ΔΔCt (Applied Biosystems User Bulletin no. 2: Rev B “Relative Quantitation of Gene Expression”).

### Protein preparation and Western blot analysis

Total protein was lysed by cold RIPA buffer. Then, a BCA assay kit (Thermo Fisher Scientific, Waltham, MA, U.S.A) measured the protein concentration of the supernatant. For Western blot experiments, IL-22 antibody (1:800, AF782, R&D Systems; Minneapolis, MN, USA), IL-22RA1 antibody (1:800, ab5984, Abcam, Cambridge, United Kingdom), α-SMA antibody (1:1000, ab28052, Abcam, Cambridge, United Kingdom), STAT3 antibody (1:800, ab68153, Abcam, Cambridge, United Kingdom) and p-STAT3 antibody (1:800, ab267373, Abcam, Cambridge, United Kingdom) was used to detect certain protein respectively, while β-actin antibody (1:5000, ab8227, Abcam, Cambridge, United Kingdom) was used as a 1:5,000 dilution to determine β-actin protein abundance. Next the membrane was incubated with HRP-conjugated goat anti- rabbit or goat anti- mouse secondary antibodies. ECL Ultra Western HRP kit (Thermo Fisher Scientific, Waltham, MA, U.S.A) was used to detect areas of luminescence, and the relative-staining intensity of each protein band was quantitated using ImageJ software (Ver. 6.0, Media Cybernetics, Inc., Rockville, MD, USA). A ratio of IL-22, IL-22RA1, STAT3, p-STAT3 and α-SMA protein intensity over β-actin protein intensity was used to quantify protein expression.

### Tissue histological observation and immunohistochemistry staining

Tissues fixed in 10% formalin were embedded in paraffin and sectioned. Eight-micrometer-thick sections were cut and stained. Briefly, IL-22RA1 or α-SMA was incubated with tissue sections at a 1:200 dilution overnight. The tissue slides were subsequently incubated with secondary antiserum (Alexa Fluor 647 goat anti-rabbit IgG; Alexa Fluor 567 goat anti-mouse IgG Molecular F Probes, Eugene, OR, USA) at 1:400 dilution.

Immunofluorescent staining of fibroblasts cells was performed on micro-coverslips in 12-well tissue culture plates. Cells were serum-starved for 24 hours and then treated with patients’ serum medium or 20 ng/mL recombinant IL-22 for 24h. Cells were washed and fixed in 4% formaldehyde, then stained with IL-22RA1 or α-SMA following incubated with required secondary antibodies. Their nuclei were stained with DAPI. Image acquisition was performed using a Zeiss confocal microscope or an upright Zeiss microscope with Axiovision software.

### Statistical analyses

All experiments were performed in triplicate, and the data were analyzed using SPSS Software (Version 23.0, IBM Corp, Armonk, NY, USA). Results are presented as the mean ± SD. Differences between groups were analyzed by using the t-test or ANOVA followed by the Tukey test. For all statistical tests, P-values < 0.05 were regarded as statistically significant

## Acknowledgments

We wish to acknowledge the excellent facility of Aier Eye Institute. Opinions, interpretations.

## References

Arshad, T., Mansur, F., Palek, R., Manzoor, S., and Liska, V. (2020). A Double Edged Sword Role of Interleukin-22 in Wound Healing and Tissue Regeneration. Front Immunol 11, 2148.

Caprioli, J., and Varma, R. (2011). Intraocular pressure: modulation as treatment for glaucoma. Am J Ophthalmol 152, 340–344 e342.

Dong, Y., Hu, C., Huang, C., Gao, J., Niu, W., Wang, D., Wang, Y., and Niu, C. (2021). Interleukin-22 Plays a Protective Role by Regulating the JAK2-STAT3 Pathway to Improve Inflammation, Oxidative Stress, and Neuronal Apoptosis following Cerebral Ischemia-Reperfusion Injury. Mediators Inflamm 2021, 6621296.

Gajda-Derylo, B., Stahnke, T., Struckmann, S., Warsow, G., Birke, K., Birke, M.T., Hohberger, B., Rejdak, R., Fuellen, G., and Junemann, A.G. (2019). Comparison of cytokine/chemokine levels in aqueous humor of primary open-angle glaucoma patients with positive or negative outcome following trabeculectomy. Biosci Rep 39.

Ji, Y.W., Mittal, S.K., Hwang, H.S., Chang, E.J., Lee, J.H., Seo, Y., Yeo, A., Noh, H., Lee, H.S., Chauhan, S.K., et al. (2017). Lacrimal gland-derived IL-22 regulates IL-17-mediated ocular mucosal inflammation. Mucosal Immunol 10, 1202–1210.

Kang, J.M., and Tanna, A.P. (2021). Glaucoma. Med Clin North Am 105, 493–510.

Kasahara, M., and Shoji, N. (2021). Effectiveness and limitations of minimally invasive glaucoma surgery targeting Schlemm’s canal. Jpn J Ophthalmol 65, 6–22.

Keller, K.E., Yang, Y.F., Sun, Y.Y., Sykes, R., Gaudette, N.D., Samples, J.R., Acott, T.S., and Wirtz, M.K. (2014). Interleukin-20 receptor expression in the trabecular meshwork and its implication in glaucoma. J Ocul Pharmacol Ther 30, 267–276.

Laroche, D., Nkrumah, G., and Ng, C. (2019). Real-World Retrospective Consecutive Study of Ab Interno XEN 45 Gel Stent Implant with Mitomycin C in Black and Afro-Latino Patients with Glaucoma: 40% Required Secondary Glaucoma Surgery at 1 Year. Middle East Afr J Ophthalmol 26, 229–234.

McGee, H.M., Schmidt, B.A., Booth, C.J., Yancopoulos, G.D., Valenzuela, D.M., Murphy, A.J., Stevens, S., Flavell, R.A., and Horsley, V. (2013). IL-22 promotes fibroblast-mediated wound repair in the skin. J Invest Dermatol 133, 1321–1329.

Saxton, R.A., Henneberg, L.T., Calafiore, M., Su, L., Jude, K.M., Hanash, A.M., and Garcia, K.C. (2021). The tissue protective functions of interleukin-22 can be decoupled from pro-inflammatory actions through structure-based design. Immunity 54, 660–672 e669.

Tham, Y.C., Li, X., Wong, T.Y., Quigley, H.A., Aung, T., and Cheng, C.Y. (2014). Global prevalence of glaucoma and projections of glaucoma burden through 2040: a systematic review and meta-analysis. Ophthalmology 121, 2081–2090.

Wolk, K., and Sabat, R. (2006). Interleukin-22: a novel T-and NK-cell derived cytokine that regulates the biology of tissue cells. Cytokine Growth Factor Rev 17, 367–380.

Yoon, C.H., Lee, D., Jeong, H.J., Ryu, J.S., and Kim, M.K. (2018). Distribution of Interleukin-22-secreting Immune Cells in Conjunctival Associated Lymphoid Tissue. Korean J Ophthalmol 32, 147–153.

Zada, M., Pattamatta, U., and White, A. (2018). Modulation of Fibroblasts in Conjunctival Wound Healing. Ophthalmology 125, 179–192.

Zhao, Y., Zhang, F., Pan, Z., Luo, H., Liu, K., and Duan, X. (2019). LncRNA NR_003923 promotes cell proliferation, migration, fibrosis, and autophagy via the miR-760/miR-215-3p/IL22RA1 axis in human Tenon’s capsule fibroblasts. Cell Death Dis 10, 594.

Zheng, Y., Danilenko, D.M., Valdez, P., Kasman, I., Eastham-Anderson, J., Wu, J., and Ouyang, W. (2007). Interleukin-22, a T(H)17 cytokine, mediates IL-23-induced dermal inflammation and acanthosis. Nature 445, 648–651.

